# Biomimetic computations improve neural network robustness

**DOI:** 10.1101/2023.10.26.564127

**Authors:** Linnea Evanson, Maksim Lavrov, Iakov Kharitonov, Sihao Lu, Andriy S. Kozlov

## Abstract

Object recognition by natural and artificial sensory systems requires a combination of selectivity and invariance. Both natural and artificial neural networks achieve selectivity and invariance by propagating sensory information though layers of neurons organised in a functional hierarchy. Both employ computational units performing AND-like operations for selectivity and OR-like operations for invariance. However, while biological neurons are intrinsically capable of switching between these operations, their artificial counterparts are hard-wired to perform only one of them. We wanted to test whether the flexible mapping between neurons and computations observed in biological neural networks is compatible with, or perhaps even useful to, artificial neural networks. To answer this question, we have developed a deep learning layer in which both selectivity and invariance operations can be performed by the same neurons. As with biological neurons, the choice of which operation an artificial neuron performs on a given input can be governed by the input strength. This flexible layer successfully outputs a combination of the two operations and, surprisingly, confers additional robustness to adversarial examples, which are inputs deliberately crafted to promote misclassification. The flexible mapping also improves accuracy when the training dataset is small, as well as when data are corrupted by certain types of noise. These results narrow the gap between biological and artificial neural networks and add a new bio-inspired approach to the arsenal of defenses against adversarial examples, which are known threats to model-based optimization and network security.

**Author summary:** The biophysical properties of a biological neuron enable it to perform both OR-like and AND-like operations over its inputs. In contrast, artificial neural networks use units that always perform only one specific operation. We wondered whether the flexibility observed in individual biological neurons could be incorporated into artificial neural networks without breaking them. If artificial neural networks required their units to perform single operations, then this requirement would either point to a fundamental difference between biological and artificial neural networks or indicate that biological neurons are somehow constrained to perform only one type of operations during object recognition. If, on the other hand, artificial units can flexibly switch between different operations, then such a result would indicate that biological and artificial neural networks are more similar than previously thought and would be an important step towards biomimetic AI. To find out, we introduced a new computational structure we call a flexible layer, in which individual units can switch between operations according to a rule (e.g., depending on input strength, or even randomly). We found that inserting the flexible layer in one or several positions in different artificial neural networks trained on several benchmark datasets not only preserves their accuracy but also makes them more robust to various perturbations and improves learning when training data is scarce.

## Introduction

After the field of Artificial Intelligence (AI) was born in the 20th century and survived the “AI winter” of the 1990s, it has rapidly diverted from its roots in modelling the brain into a myriad of sub-fields such as machine learning, deep learning, reinforcement learning and others. These are now standalone scientific fields with their own methodologies and enormous amounts of accumulated knowledge, but with fewer and weaker direct links to the field of neuroscience [1]. Yet opportunities to establish such links and benefit from them abound today thanks to the rapid development of experimental and theoretical neuroscience [2].

Neuroscience and AI are historically intertwined, with multiple object-recognition models (e.g., HMAX) inspired by the anatomy and physiology of the visual cortex [2, 3, 4]. For instance, Hubel & Wiesel discovered two different types of visual cortical cells which they named simple and complex cells [5]. These cell types display selectivity to stimuli of different spatial orientations and, at the level of complex cells, invariance to the stimulus position within the receptive field. Many years later, state-of-the-art object recognition models such as Convolutional Neural Networks (CNNs) that only distantly mimic the structure of the brain, also use feature recombination functions to achieve selectivity and invariance. In biological systems, selectivity is classically defined in terms of the optimal stimulus for a neuron [6]. For instance, some neurons of the visual cortex are selective to particular orientations of bars in space or more complex features, such as entire objects. Invariance in object recognition is defined as the ability to correctly classify visual objects in their previously learned object-name category, despite variations in object appearance due to object-identity preserving transformations, such as changes in object and viewer position, illumination conditions, and occlusion by other objects [7].

A basic neural operation corresponding to selectivity consists of taking dot products between input vectors and a set of filters whose values can be modified through learning. In a Convolutional Neural Network (CNN), selectivity is achieved by the convolutional layer units. At the same time, common invariance functions in hierarchical models including CNNs are pooling or aggregation functions, such as MAX-pooling [8]. CNNs and other models, however, assume a fixed neuron-to-computation mapping, where each unit performs only one specific operation. Flexible neuron-to-computation mapping was demonstrated previously in the auditory system of songbirds and was shown to be dependent on the strength of inputs received by neurons [9]. Since the flexible mapping is rooted in basic neuronal biophysics and therefore likely to be a general property of various sensory modalities [10], we explore whether hierarchical object-recognition models, in particular CNNs, can benefit from having a similar property.

### Our Approach

We propose a new type of CNN layer, a Flexible Layer, whose units switch according to a specified rule (e.g., in an input-dependent manner) between performing convolution or pooling operations during the forward pass, both during training and inference. Namely, the strength of the signal coming into the Flexible Layer unit defines whether that unit performs a convolution or pooling operation. Similar to dropout, this method creates sub-networks, but there are two key differences: 1) the sub-networks are not strictly sub-sampled, because the units are not omitted, but are filled with a different operation result and some information is retained; 2) instead of dropping out the units based on probability there is a decision rule that governs when a specific operation is performed. This decision rule can be, for example, threshold-based. Figure 1 shows a schematic representation of the proposed architecture which uses a threshold on the input signal strength to switch between performing a convolution operation and a MAX-pooling operation. Thus, such a Flexible Layer possesses a new type of parameter (shown as *T*), which is, in principle, learnable and defines the threshold at the level of a single unit (neuron).

**Fig 1.**
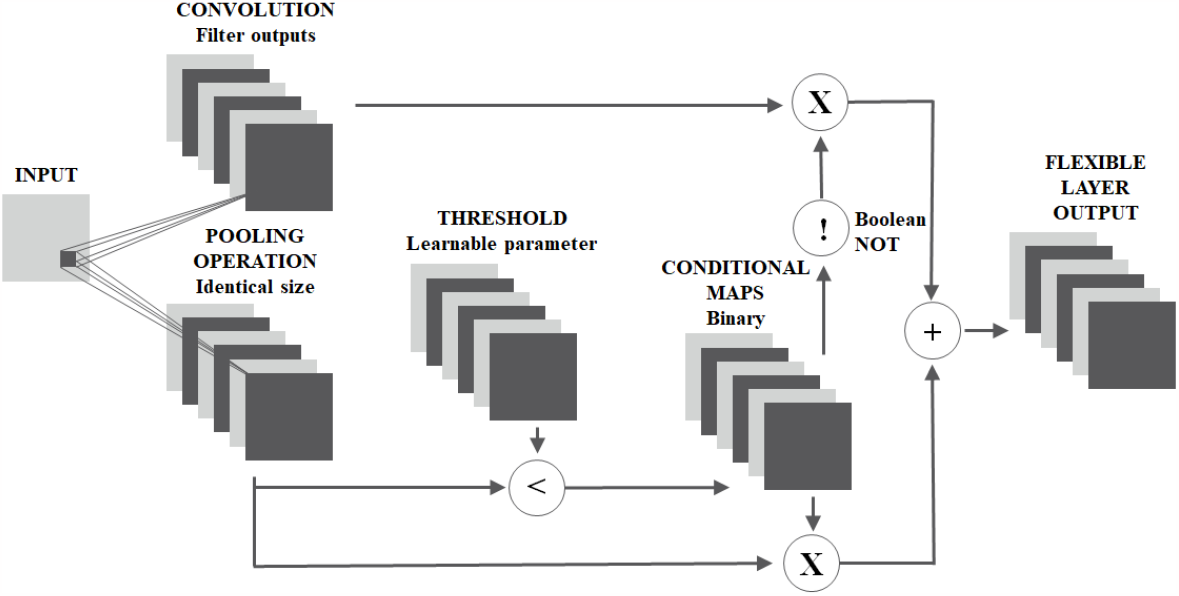
Schematic representation of the Flexible Layer input-output relationship. For each input image, convolutional filtering is applied, and in addition MAX-pooling is performed in parallel. The result of the MAX-pooling operation is compared with a threshold *T* to create a conditional mask. The mask is used to select the units for which the result of the MAX-pooling operation will be used in the layer output. To select which units will use the convolutional output, we use the logical complement of the conditional mask.

Recent advancements in CNNs have led to performance at a level equal to or above that achievable by humans in object recognition. Nonetheless, modern deep learning architectures have been shown to be susceptible to adversarial attacks [11, 12, 13], which are based on adversarial examples. Adversarial examples are input perturbations that are found by optimizing the input to maximize the prediction error and are generally (but not necessarily) visually undetectable by humans [14]. One can distinguish white-box attacks, which assume complete knowledge of the model under attack, and black-box attacks, which assume knowledge of only the model’s input-output characteristics.

Some work has also been done on transferability of black-box attacks that can fool multiple machine learning models and demonstrated that it can also fool time-limited humans [15]. These findings indicate a promising avenue of study on whether computational mechanisms deployed by the brain might contribute to adversarial attack robustness and can be translated to better object recognition models. Similarly to how dropout was inspired by the mixability theory in evolutionary biology [16] and proposed in machine learning to reduce overfitting [17] along with defensive dropout mechanism to protect against adversarial attacks [18], we suggest flexible mapping architecture as a possible bio-inspired defence mechanism for CNNs.

## Materials and methods

### Flexible Layer Structure

The Flexible Layer outputs a combination of the output of a convolutional layer and MAX-pooling layer computed in parallel (Fig. 1). The specific selection between convolution and MAX-pooling on the element-level is decided by an element-wise comparison between the MAX-pooling output and a threshold tensor, *T*. This yields a boolean mask which can be applied to select the units for which the MAX-pooling result will be chosen. Similarly, the logical complement of this mask is used to select the units for which the convolution result will be used. The final output is thus a combination of both convolutional filtering and MAX-pooling. This is called the standard Flexible Layer, and is used in all Flexible Networks unless specified otherwise.

We define two additional thresholding schemes. The first one includes fixed, uniform thresholds for the threshold tensor, *T*. These values are defined during network initialisation and are not updated during the training procedure. The second scheme uses a threshold tensor, *T*, with random values. These values randomly change between 0 and 1 with equal probability at each forward pass during both training and inference. Similarly, the parameters for the threshold assignment in this case are not adjusted during the training process.

### Operation Selection Implementation

All operations in a neural network are required to be differentiable to maintain the ability to perform backpropagation in order to calculate the gradients used to learn the network parameters. With this in mind, we implement the boolean less-than (*<*) operation and subsequent conditional masking as a sum of sigmoidal functions, where each sigmoidal function acts as a pseudo-binary factor to either input (either the convolution or max pooling output). Mathematically, this can be described as:

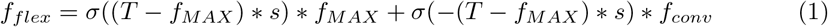

where *f*_*flex*_ is the output of the Flexible Layer, *σ*() is the sigmoidal function, *T* is the threshold tensor, *s* is a scaling factor, *f*_*MAX*_ is the output for the MAX-pooling operation and *f*_*conv*_ is the output of convolution on the input. The scaling factor *s* is a constant to drive the input into the saturation region of the sigmoid. In our implementation, *s* was chosen to be 50.

### Network Architectures

The Flexible Layer was incorporated into three different baseline architectures to explore its effects on each model’s performance. The specific location of the Flexible Layer within each network is shown in Figure 2. The main tests were performed on the VGG16 network, where we replaced the second convolutional layer in the first block (Fig. 2A). We call this architecture the Single Flexible Layer VGG16 network. We define the baseline network as this same network with the Flexible Layer inserted, but with the threshold tensor set such that convolution is chosen for all inputs, thereby making the resulting output equal to that of a regular convolution layer. We train the VGG16 networks on the CIFAR10 dataset [19] for the adversarial attack tests, and on Imagenette [20] for the frequency response analysis.

**Fig 2.**
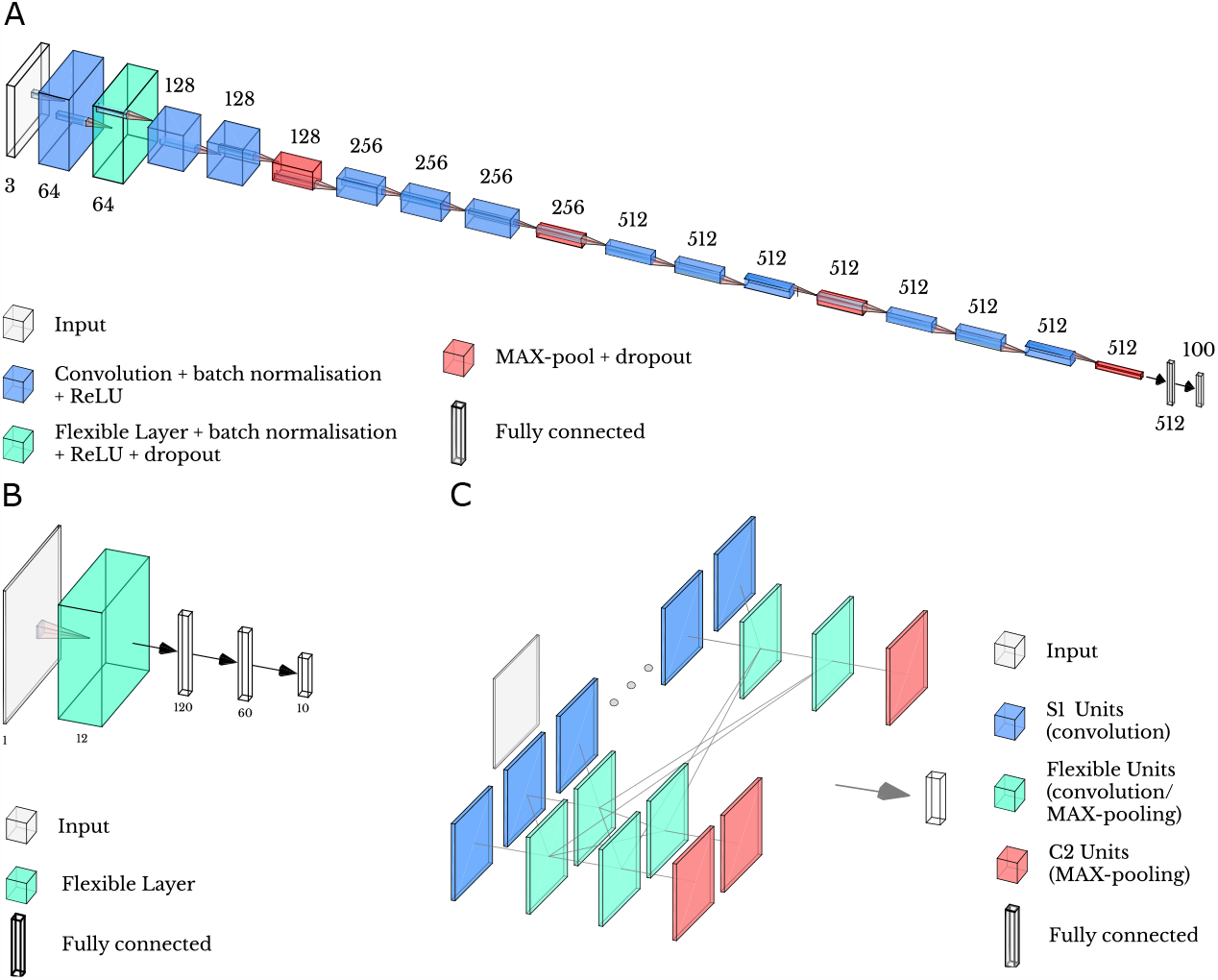
Schematics for incorporating the Flexible Layer into existing architectures. (**A**) Schematic of the VGG16 architecture with the Flexible Layer inserted. Numbers indicate depth of the tensor at each step.(**B**) Schematic of the architecture used for the FMNIST dataset. (**C**) Schematic for the HMAX model with Flexible Layers instead of the C1 and S2 units. There are 7 such units in the HMAX model.

In addition, we train a smaller network (Fig. 2B) with a Flexible Layer on the FMNIST dataset [21]. Again we compare the performance against a baseline where the threshold is inactivated, so that only convolution is returned from the layer.

We also insert the Flexible Layer into the HMAX model (Fig. 2C). The HMAX model of object recognition in the primate ventral stream is historically a very important model that sits at the intersection between Hubel & Wiesel’s neurophysiological findings and more modern deep neural networks that are built around the same underlying principle, namely hierarchies of units dedicated to building selectivity and invariance. In the standard HMAX model, complex-like C1 cells perform MAX-pooling with different size kernels, and simple-like S2 cells take the Euclidian distance between C1 layer output and a convolutional filter that is learned during training [22, 23]. In our implementation, the Flexible HMAX has Flexible Layers inserted in the C1 and S2 layers and, in order to ensure any improvement in accuracy is not due solely to adding convolutional layers into the model, a Convolutional Baseline with Flexible Layers executing convolution in the C1 and S2 layers was trained for comparison. Here convolution is performed on inputs above the threshold and MAX-pooling on inputs below. A model without any modifications is also trained for comparison; we call this “Baseline HMAX”. The HMAX networks were trained on the FMNIST dataset. Additional details of network architectures and training datasets can be found in the Supplementary Information.

### Adversarial Attacks

#### FGSM Attacks

One method of attack is the Fast Gradient Sign Method (FGSM), in which the gradient of the network is not known to the adversarial attacker; it simply adds noise of magnitude *ϵ* to the images in the direction which would increase the cost function, to create perturbed images **x**_*p*_ according to the following equation [24, 25]:

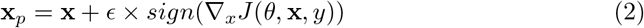

where **x** are the input images with target labels *y*, to a network with cost function *J* and hyperparameters *θ*. We run FGSM attacks with perturbations *ϵ* of sizes from 0.05 to 1. Accuracy for one attack is reported on the whole validation dataset of 10,000 examples and average accuracy is calculated from 10 iterations of each attack. We follow examples provided by the PyTorch FGSM Tutorial [26].

#### PGD Attacks

Projected Gradient Descent (PGD) attacks use the gradient of a network to find the direction in which to most efficiently add noise to cause misclassification [27]. PGD attacks come in both white- and black-box varieties. White box attacks have access to the model and all its parameters, while black-box attacks use an estimation of the model, which is often a simplified network. Attacks are made iteratively: first, noise is added randomly, then that perturbed image is passed through the network and the direction of the loss gradient is calculated. Then a “step” is taken in the attack: noise is added in the direction of the gradient to maximise loss. The noise is added with a certain maximum perturbation magnitude *ϵ*.

We run white- and black-box PGD attacks on the Single Flexible Layer VGG16 networks. The attacks are run 100 times and averaged accuracies are presented. For both white- and black-box PGD attacks the settings are: *ϵ* = 0.031, with 20 steps and step size of 0.003, tested on the whole validation dataset. For the black-box PGD attacks we use the “Conv2D model” [27] which has 4 convolutional, 2 max pooling and 2 linear layers, to estimate the network gradient. The details of that model can be found in Supplementary Information: Network Definitions. Code is based on the work by Zhang et al.[28] and Yu [29].

#### Real-world corruptions

Hendrycks and Dietterich [30] presented a dataset of 19 image corruptions relevant to natural visual scenes. These corruptions are split into 5 categories: weather, digital, noise, blur, other (Table 1). Five levels of corruption severity are available. They were handcrafted by humans, e.g., in high severity snow corruption, the density of snow is visually higher and overall visibility is poorer than it is for low severity levels. First, the Flexible and Baseline Networks were trained on the original unaltered 10 class ImageNet data. These corruptions were then applied to the validation set and the inference accuracy on this set of images was recorded.

**Table 1.** Types of real-world corruptions applied to the validation dataset, developed by Hendrycks and Dietterich [30]. Each corruption type has five levels of severity.

| blur | digital | noise | weather | other |
| --- | --- | --- | --- | --- |
| defocus blur | contrast | gaussian noise | snow | speckle noise |
| glass blur | elastic transform | shot noise | frost | gaussian blur |
| motion blur | pixelate | impulse noise | fog | spatter |
| zoom blur | jpeg compression |  | brightness | saturate |

#### Spectral analysis

To characterise the frequency responses of our different architectures, we considered the classification of images with limited spectral content. Networks were trained on ImageNette data. For inference, the validation set was converted to Fourier space and a Gaussian bandpass filter was applied to each image in bands of 5 Hz, between 1 and 160 Hz (as 160 Hz is roughly the maximum possible frequency or maximum Euclidean distance in Fourier space for 224x224 images). Then, the inverse Fourier transform was applied to the bandpassed images to return to RGB space. The inference accuracy of the Baseline and Flexible VGG16 models was recorded for each of these bands.

#### Code availability

PyTorch code for the Flexible Layer and example notebooks for the analyses are available online: https://gitlab.com/kozlovlabcode/flexible-networks

## Results

### Flexible computation and network accuracy

First we test the effect of the Flexible Layer when inserted into a simple small convolutional network (Fig. 2B). We observe that introducing flexible mapping leads to validation accuracy that is 2-6% lower compared to the baseline network (Table 2).

**Table 2.** Comparison of validation accuracy after training, and FGSM attack accuracy between different implementations of thresholds in the small network. Uniform and standard thresholds are defined in Methods. The standard reverse threshold performs MAX-pooling above threshold instead of convolution. We use a simple single-shot FGSM attack with ϵ **= 0.1**, and report the accuracy on images in the validation set. We trained 10 different initialisations of each network and report the mean accuracy of all the models and the corresponding standard error of the mean (SEM) with and without FGSM attack.

| Architecture Description | Validation Accuracy (%) | SEM (%) | FGSM (%) | SEM (%) |
| --- | --- | --- | --- | --- |
| Baseline Network | 87.7 | 0.54 | 27.3 | 0.89 |
| Uniform Threshold Network | 86.3 | 0.50 | 44.4 | 0.82 |
| Standard Threshold Network | 85.2 | 0.53 | 36.4 | 0.85 |
| Standard Reverse Threshold Network | 84.6 | 0.93 | 31.9 | 1.12 |

To test whether the network optimises around the values of a threshold, we train several of these small flexible networks (Fig. 2B) with fixed, uniform thresholds. We then test these trained networks with different fixed uniform threshold values set at the time of inference to see whether they have become optimised for their given *T* values. The results confirm that the network optimises around the values of the threshold, and performs worse if a different fixed uniform threshold is used for validation (Fig. 3). Second, after inserting the Flexible Layer into the HMAX network, the Flexible HMAX outperforms both the Baseline and Convolutional Baseline HMAX networks by 20% and 6% test accuracy respectively after 150 epochs (Fig. 4A). To explore the effect of the Flexible Layer in a more complex architecture, we add it to the VGG16 network. The Flexible Layer does not compromise the validation accuracy compared to the baseline in all of its configurations (Fig. 4B).

**Fig 3.**
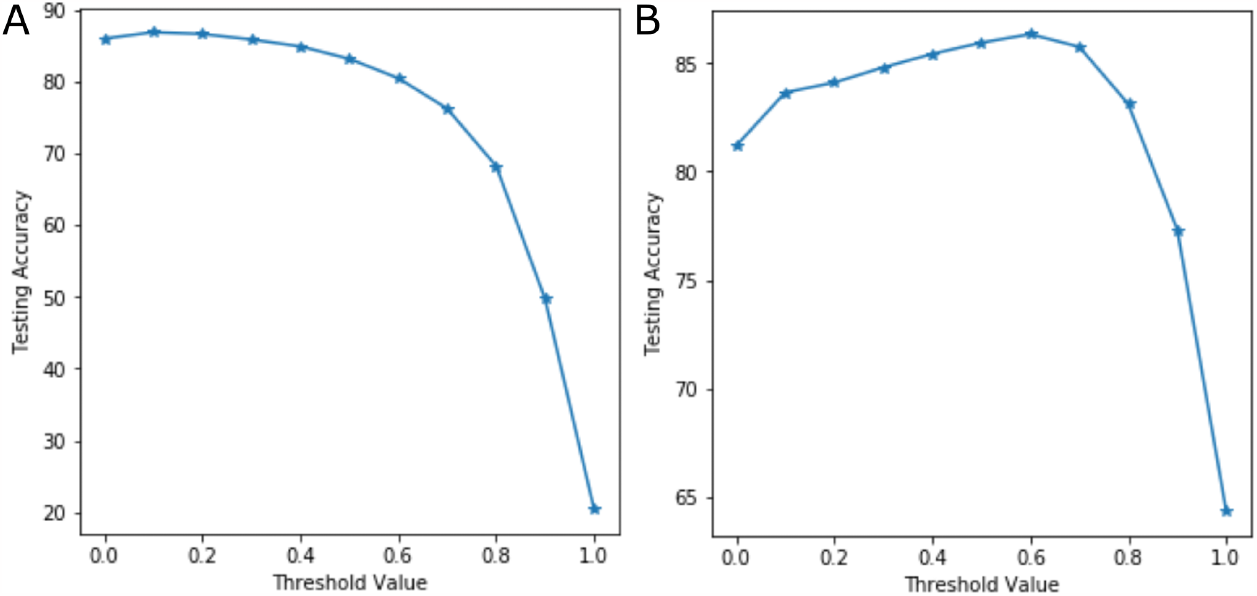
Validation accuracy of a small flexible layer network (Fig. 2B) trained with fixed uniform thresholds. (**A**) Small flexible layer network trained with a fixed uniform threshold of 0.1. (**B**) Small flexible layer network trained with a fixed uniform threshold of 0.6. Fixed uniform threshold was varied at the inference stage during validation.

**Fig 4.**
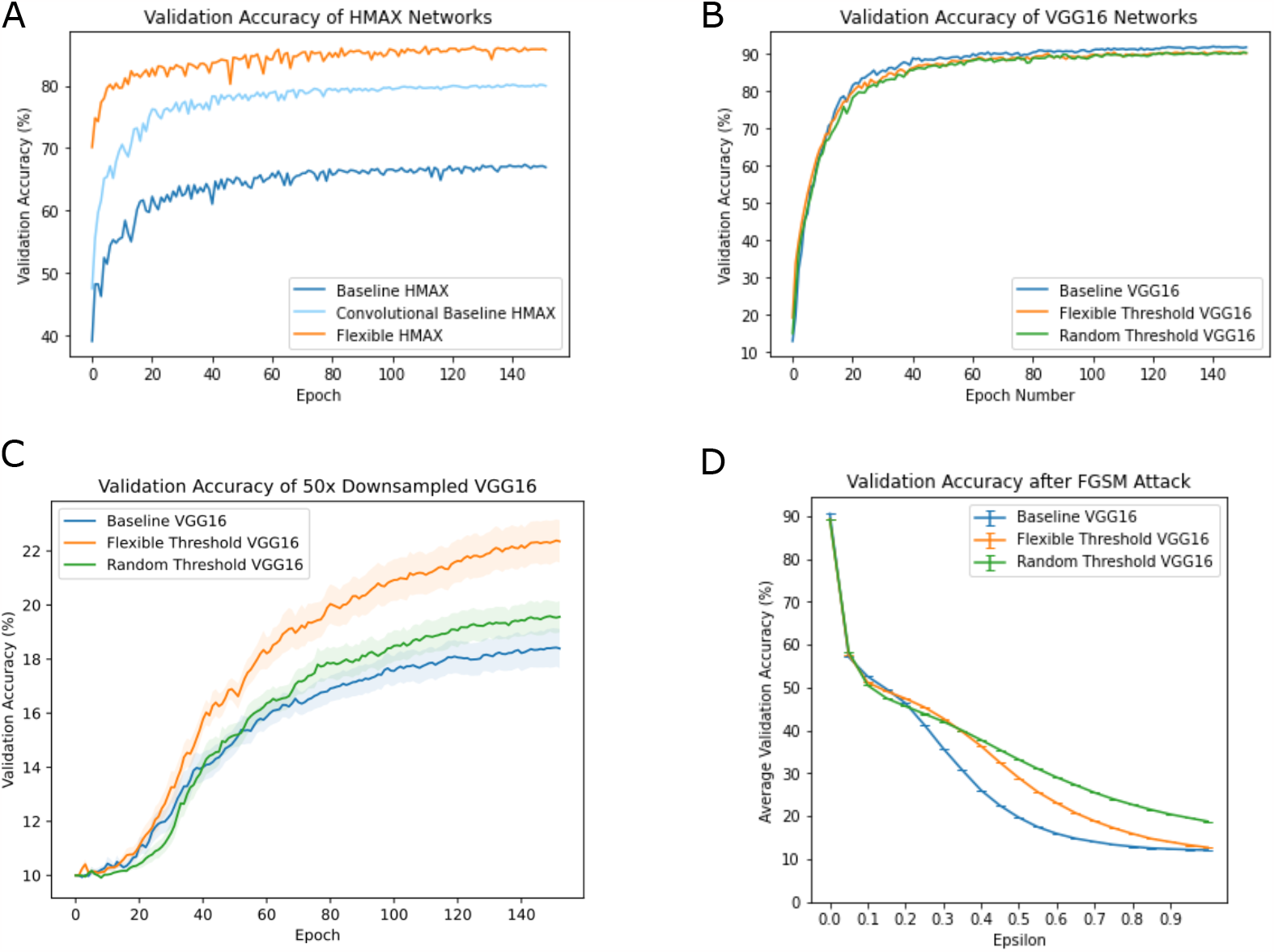
Comparison of performance between Baseline and Flexible Layer networks. (**A**) HMAX models [22] trained on the FMNIST dataset, the Flexible HMAX has Flexible Layers inserted in the S2 and C1 layers, the Convolutional Baseline has convolutional layers in the S2 and C1 layers. (**B**) Validation accuracy of the different VGG16 networks trained on CIFAR10. The Baseline VGG16 network (blue), the Single Layer Flexible network with trained threshold values (orange) and the Flexible network with randomly chosen threshold values (green). (**C**) A VGG16 model with a single Flexible Layer in Block 1, whether the threshold is learned or random shows increased validation accuracy during training on the CIFAR10 dataset that has been downsampled 50 times. Shading is standard error of the mean across 10 networks. (**D**) Average validation accuracy of VGG16 Networks trained on the CIFAR10 dataset after FGSM attacks. Accuracy for one attack is reported on the whole validation dataset of 10 000 examples and average accuracy is calculated from 100 iterations of each attack. *ϵ* = 0 is the same as no attack. Error bars are standard error of the mean.

Lastly, we show the benefit conferred by the Flexible Layer on training with small amounts of data. We trained 10 VGG16 networks on datasets of approximately 1000 images. The datasets were the CIFAR10 dataset downsampled 50 times, with images sampled evenly from each of the 10 classes. We show that the Flexible Layer performs better than the baseline network during all the epochs of the training procedure (Fig. 4C), and regardless of random seed used to generate the training data.

Overall, these results show that the bio-inspired, flexible operations can be successfully incorporated into various artificial neural networks. In general, they do not compromise network accuracy and can even lead to an improvement compared to the baseline, especially when training data is scarce.

### Flexible computation confers robustness against a range of perturbations

Adversarial robustness was tested on the small convolutional network (Fig. 2B) and the VGG16 network. The adversarial robustness of the small network was tested by employing an FGSM attack with *ϵ* = 0.1. We observe that the robustness against FGSM attack is 6-15% greater than that of the baseline model (Table 2). This effect weakens when we reverse the operation selection of the Flexible Layer.

For the VGG16 network, adding a Flexible Layer led to an increase in validation accuracy compared to the Baseline under a simple FGSM attack for disturbance length *ϵ >* 0.22 (Fig. 4D). For the PGD attacks, adding a Flexible Layer increased test accuracy, relative to the Baseline network, after the white-box attack (Table 3) by 8.76%. The test accuracy after the relatively weaker black-box PGD showed little difference in comparison with the Baseline network; in both cases the accuracy was greater than after the white-box attack, as expected.

**Table 3.** Comparison of PGD classification accuracies between the Baseline VGG16 model and the Flexible VGG16 model trained on CIFAR10. Accuracies for both the white- and black-box configurations are presented.

| Model | Natural | PGD white-box | PGD black-box |
| --- | --- | --- | --- |
| Baseline VGG16 | 91.42% | 24.81% | 60.21% |
| Flexible VGG16 | 90.18% | 33.57% | 60.41% |

Given the existence of flexible computation in biological systems, we sought to determine whether this computational principle would confer added robustness to naturally occurring visual perturbations by testing our flexible network performance on a natural perturbation benchmark [30]. As illustrated by Fig. 5A, Single Layer Flexible VGG16 shows similar or marginally higher robustness accuracy over the 5 corruption categories of the Real-World Corruptions dataset, compared to Baseline VGG16. All Layer Flexible VGG16 is the least robust for “blur”, “digital” and “weather”, but the most robust for “noise” (7% over Baseline at highest severity).

**Fig 5.**
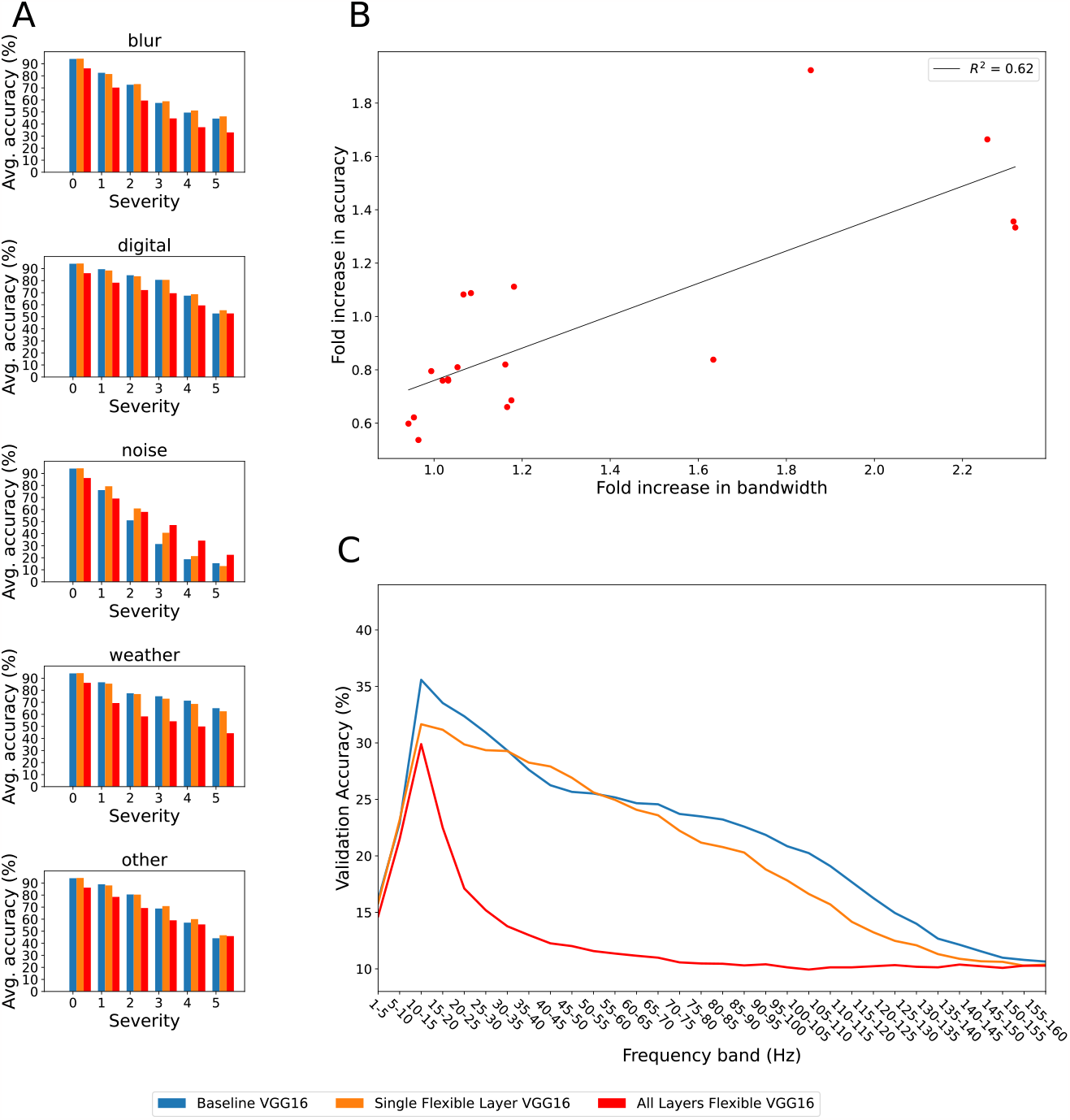
Performance on real-world corruptions dataset and spectral analysis of Baseline and Flexible Layer networks. **(A)** Validation accuracy of Baseline, Single Layer Flexible and All Layers Flexible VGG16 networks on the real-world corruptions dataset. Accuracy is averaged over all ImageNet classes and categories of corruptions (blur, digital, noise, weather and other, as seen in Table 1). Six levels of severity are shown for each of the five corruption categories, where severity = 0 represents uncorrupted images. **(B)** Increasing bandwidth, by corrupting the images, results in increasing validation accuracy of the All Layer Flexible VGG16 network relative to the Baseline VGG16 network. Relative accuracy is denoted as the fold increase in classification accuracy of the All Layer Flexible VGG16 over Baseline VGG16 at corruption severity 5. Fold increase in bandwidth is quantified as the ratio of the full width at half maximum (FWHM) value of the spatial Fourier decomposition at corruption severity 5 to the FHWM of the uncorrupted image. This ratio is averaged over every image within each of the corruption categories. An increase in bandwidth represents stronger high frequency content in an image. **(C)** Validation accuracy of Baseline, Single Flexible Layer and All Layers Flexible VGG16 models on ImageNet with respect to the center frequency of a Gaussian bandpass filter applied to the input images. Bandwidth of Gaussian bandpass filters was set to 5 Hz.

We wondered whether the robustness of Flexible Networks to adversarial attacks and noise, under certain conditions, could be due to the effect of the Flexible mapping causing the network to rely more on low-frequency information in the images, rather than high-frequency information, making it more similar to humans who are not perturbed by the addition of high-frequency noise [24]. To test this idea, we performed a spectral analysis of the Flexible and Baseline networks on validation images from ImageNet and found a positive correlation between the increase in bandwidth of corrupted images and the increase of robustness accuracy of Flexible Networks relative to Baseline (Fig. 5B). All corruptions alter spectral content of images, but “shot noise”, “gaussian noise”, “impulse noise” from the “noise” category and “speckle noise” from the “other” category (see Table 1) produce the biggest increases in high frequency content (top right of Fig. 5B).

Single Flexible Layer network has a similar response to Baseline when low-frequency image content is removed. Accuracy of All Layers Flexible network drops sharply when 5-20 Hz information is absent, as shown in Fig. 5C, indicating that it indeed relies on this low-frequency information.

We hypothesised that this improved performance on lower frequency components within images implies that classification outcome is largely driven by general shapes of objects rather than fine textural details. This in turn would imply that the network would generalise better to out-of-distribution examples. We tested the flexible network on data generated by Evans, Malhotra and Bowers [31]. This dataset stylised images derived from the CIFAR-10 dataset. The stylised images are categorised as “contours”,”line drawings”, “silhouettes”. As the images were 224x224, the standard 32x32 training set of CIFAR-10 had to be upscaled using Lanczos resampling, following the original methodology [31]. All Layers Flexible VGG16 indeed achieved better generalisation by 6%, 14% and 7% in “contours”, “line drawings” and “silhouettes”, respectively (Fig. 6).

**Fig 6.**
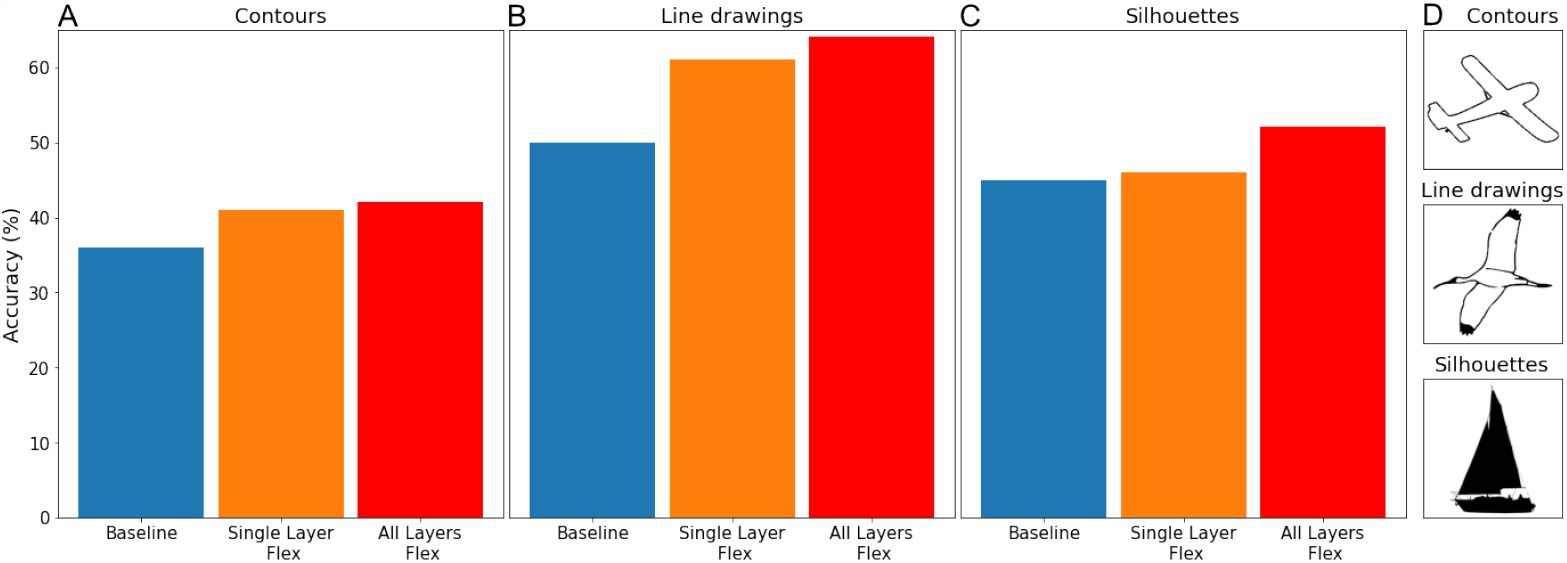
Out-of-distribution performance of Flexible Layer networks. Classification accuracy on out-of-distribution images of Baseline models versus Flexible VGG16 models. Classification accuracy of Baseline and Flexible Layer networks on o.o.d images (contours (**A**), line drawings (**B**) and silhouettes (**C**)). (**D**) Examples of the different types of o.o.d. images of each of the three categories.

## Discussion

Deep learning has its roots in basic neuroscience, and this study was motivated by a basic neuroscience question. Historically, since Hubel & Wiesel’s work [5], the dichotomy between simple and complex cells led to a commonly held view about how selectivity and invariance are achieved in biological and artificial neural networks, as well as to specific implementations in various biologically-inspired artificial neural networks.

Every biological neuron receives synaptic input, and every spiking neuron converts this input into a firing rate. The input-output relationship between the synaptic current and the firing rate is non-linear. The action-potential threshold, in particular, will result in AND-like operations performed on weak inputs (inputs that individually are sub-threshold) and OR-like operations performed on strong inputs. Given that the strength of input is not fixed but can change, how can a neuron be assigned a specific operation, or a feature recombination function, and not switch between them? Perhaps a neuron does not need to be assigned a specific operation and can freely switch between them?

In principle, there are at least three possibilities. First, it is possible that, just as in the HMAX model, there must be separate AND-like and OR-like neurons implementing selectivity and invariance, respectively. In this case, it is unclear how this dichotomy can survive when synaptic weights change, e.g., during adaptation. There would have to be some mechanisms that maintain the fixed neuron-to-computation mapping. No such mechanisms are known. Attention could be one such mechanism: it would provide additional input current to otherwise adapted neurons. The operations could then “survive” in the locus of attention. This possibility was mentioned in Kozlov & Gentner [9]. Based on our results here, we feel that this scheme is too complicated and unnecessary, given the alternatives. The second possibility is that neurons perform one or the other operation depending on the strength of the input. For example, they could perform an AND-like operation when inputs are sufficiently weak and an OR-like operation when they are strong. In this case, when a (non-adapted) neuron is presented with its preferred stimulus, it will receive a strong input and will therefore perform an OR-like computation, for example a MAX, thus achieving some invariance or resistance to corruptions of this preferred stimulus. This scenario is supported by well-understood biophysics and can be implemented by the same canonical circuit [10]. The mapping between neurons and computations would be not fixed, but flexible. Our results here show that a neural network with such a binary mask works as well as, or even better than the same network but without the flexible mapping.

The third possibility is that no specific mapping between individual neurons and computations is required, and each neuron is free to randomly “choose” its instantaneous feature recombination function based on its intrinsic excitability and the overall excitation-inhibition balance. This “choice”, however, is inconsequential, provided the population of such neurons as a whole performs operations for selectivity and invariance. In this scenario, even a unimodal distribution of synaptic input strengths and the spiking nonlinearity can still produce an apparent functional dichotomy that can be interpreted as simple-like and complex-like cells (see [32]), but such a dichotomy would be superficial and of no real functional significance. This possibility is attractive in its simplicity. Our results with the random binary mask (random switching between the operations) show that this implementation not only works but provides an additional benefit, compared to both the baseline and the non-random flexible mapping, at greater values of input corruption (greater epsilon in Fig. 4D).

Spectral analysis has revealed that Flexible Layers focus on low-frequency content in images. In both adversarial and natural settings, uncorrelated high frequency noise tends to obscure some features that neural networks can exploit for classification. Jo & Bengio [33] highlighted the tendency of ANNs to learn superficial statistical regularities from the data, rather than higher-level abstractions. We have confirmed this bias (removal of high frequency content degrades performance, Fig. 5C) and demonstrated the ability of flexible computations to alleviate the deterioration of accuracy in response to increased noise intensity. This implies that Flexible Networks have a more robust encoding of information contained in lower frequency range, which relates to higher-level semantic concepts.

The “blocky” pattern in activation maps (likely to be associated to MAX-pooling) emerging in relatively shallow layers of the Flexible architectures suggests their tendency to extract representations which are more invariant (Fig. S1). This comes, however, with a price: networks miss out on selectively recognising high frequency textural features that could potentially yield higher validation accuracy. Nevertheless, downplayed learning of the dataset statistics is independent from generalization performance, which is increased.

Out-of-distribution generalisation is particularly illustrative. CNNs have shown to exhibit a texture-bias [34], but as noted by Malhotra, Evans, and Bowers [35], biomimetic modifications (e.g., Gabor filters in the front end) can rectify that bias. Shape-texture distinction is related to the importance of low-frequency information: Flexible Networks appear to have a stronger shape bias, quantitatively shown by 14% improvement over baseline on o.o.d. line drawings of the same classes of data (Fig. 6).

Prior studies that aimed at introducing biologically-inspired computations into CNNs reported increased adversarial robustness. Reddy et al. [36] used non-uniform sampling, mimicking non-uniform photoreceptor density in the retina, as well as spatially varying receptive-field sizes inspired by the visual cortex, and they found that such mechanisms could improve adversarial robustness with white-box PGD attacks. Dapello et al. [37] used a front-end that simulated the primary visual cortex before feeding images into CNNs, and they found an increase in both adversarial robustness under white-box PGD attacks as well as increased robustness to common image corruptions (the same ones we used in our study). Our approach is different in that we did not simulate any specific property of the visual system but instead allowed units in the flexible layer to switch between operations depending on input, or even randomly, an approach that is inspired by the basic biophysics of biological neurons.

A recent work [38], which pursued an interesting direction of neural manifold matching by ANN regularisation, corroborates our finding of low frequency preference of biomimetic models. The emergence of several links to this idea, such as blurry image training [39] and low-norm image perturbations capable of biasing human perception in a targeted way [40] hints that ANNs, in general, may be faithful models of robustness in biological visual systems. Particular approaches granting robustness might vary, but appear to have shared frequency preference properties, with the biophysically-grounded Flexible Layer fitting into this framework.

## Conclusion

Reminiscent of the broader field of biomimetics, the interplay between nature and engineering has provided AI systems with some valuable insights with regards to architecture and design. Just as aviation has initially benefited from studying avian flight mechanics, or the development of hydrophobic surfaces was inspired by the lotus leaf, the first neural networks were inspired by the organisation of the brain. By incorporating flexible recombination functions, one of our objectives is to enhance the adaptability and robustness of CNNs, better equipping them to the challenges posed by real-world problems. Specifically, we have shown that introducing flexible recombination functions observed in biological neurons into artificial neural networks improves learning when training data are scarce and increases robustness to adversarial as well as real-world perturbations.

## Acknowledgments

We thank Prof Anil Bharath for helpful discussions.

## Author Contributions

**Conceptualization:** Kozlov

**Data Curation:** Evanson, Lavrov, Lu, Kharitonov, Kozlov

**Formal Analysis:** Evanson, Lavrov, Kharitonov

**Funding Acquisition:** Kozlov

**Investigation:** Evanson, Lavrov, Kharitonov

**Methodology:** Kozlov

**Project Administration:** Kozlov

**Resources:** Kozlov

**Software:** Evanson, Lavrov, Lu, Kharitonov

**Supervision:** Kozlov, Lu

**Validation:** Evanson, Lavrov, Lu, Kharitonov, Kozlov

**Visualization:** Evanson, Lavrov, Lu, Kharitonov

**Writing – Original Draft Preparation:** Evanson, Lavrov, Lu, Kharitonov, Kozlov

**Writing – Review & Editing:** Evanson, Lu, Kharitonov, Kozlov

## Acronyms

AI: Artificial Intelligence.
ANN: Artificial Neural Network.
CNN: Convolutional Neural Network.
CNNs: Convolutional Neural Networks.
FGSM: Fast Gradient Sign Method.

## Supporting information

### Network Definitions

Layers are defined as:

Conv2d(number of input channels, number of output channels, kernel size, stride, padding)

MaxPool2d(kernel size, stride, padding)

BatchNorm2d(number of features, momentum)

Dropout2d(probability to be zeroed)

Linear(in features, out features). All linear layers learn bias.

### Single Layer Baseline/Flexible VGG16

1. Conv2d(3, 64, (3, 3), (1, 1), (1, 1))
2. BatchNorm2d(64, 0.1)
3. ReLU()
4. Flexible Layer
  a. Conv2d(64, 64, (3, 3), (1, 1), 0)
  b. MaxPool2d(3, 1, 0)
  c. Sigmoid()
5. BatchNorm2d(64, 0.1)
6. ReLU()
7. Dropout2d(0.3)
8. Conv2d(64, 128, (3, 3), (1, 1), (1, 1))
9. BatchNorm2d(128, 0.1)
10. ReLU()
11. Conv2d(128, 128, (3, 3), (1, 1),(1, 1))
12. BatchNorm2d(128, 0.1)
13. ReLU()
14. MaxPool2d(2, 2, 0)
15. Dropout2d(0.4)
16. Conv2d(128, 256, (3, 3), (1, 1), (1, 1))
17. BatchNorm2d(256, 0.1)
18. ReLU()
19. Conv2d(256, 256, (3, 3), (1, 1), (1, 1))
20. BatchNorm2d(256, 0.1)
21. ReLU()
22. Conv2d(256, 256, (3, 3), (1, 1), (1, 1))
23. BatchNorm2d(256, 0.1)
24. ReLU()
25. MaxPool2d(2, 2, 0)
26. Dropout2d(0.4)
27. Conv2d(256, 512, (3, 3), (1, 1), (1, 1))
28. BatchNorm2d(512, 0.1)
29. ReLU()
30. Conv2d(512, 512, (3, 3), (1, 1), (1, 1))
31. BatchNorm2d(512, 0.1)
32. ReLU()
33. Conv2d(512, 512, (3, 3), (1, 1), (1, 1))
34. BatchNorm2d(512, 0.1)
35. ReLU()
36. MaxPool2d(2, 2, 0)
37. Dropout2d(0.4)
38. Conv2d(512, 512, (3, 3), (1, 1), (1, 1))
39. BatchNorm2d(512, 0.1)
40. ReLU()
41. Conv2d(512, 512, (3, 3), (1, 1), (1, 1))
42. BatchNorm2d(512, 0.1)
43. ReLU()
44. Conv2d(512, 512, (3, 3), (1, 1), (1, 1))
45. BatchNorm2d(512, 0.1)
46. ReLU()
47. MaxPool2d(2, 2, 0)
48. Dropout2d(0.5)
49. Linear(512, 100)
50. Dropout(0.5)
51. BatchNorm1d(100, 0.1)
52. ReLU()
53. Dropout(0.5)
54. Linear(100, 10)

### Small Network for FMNIST experiments

1. Flexible Layer
  a. Conv2d(1, 12, (5, 5), (1, 1), 0)
  b. MaxPool2d(5, 1, 0)
  c. Sigmoid()
2. Linear(6912, 120)
3. Linear(120, 60)
4. Linear(60, 10)

#### HMAX Model

The HMAX architecture is as defined in [41, 42]. Its performance is explained as follows: The first layer, which takes in raw pixel data is composed of S1 cells, and each S1 cell is a Gabor filter of different wavelength. A Gabor filter is a bandpass filter: a sinusoid with some wavelength, multiplied by an envelope which is a Gaussian of some standard deviation. In the HMAX network the standard deviation of the Gaussian envelope scales with the wavelength, so the resulting filter detects different thicknesses of edges with those different wavelengths. There are also 4 orientations (90, -45, 0, 45) of each edge. Next, each complex-like C1 cell max pools the output of two S1 cells (each with their 4 different orientations). A different size of max pooling kernel is used for each C1 unit, which accounts for different sizes of objects in the image. Then the simple-like S2 cells take the Euclidean distance between C1 layer output and a convolutional filter which is learned during training. Finally, the C2 cells again max pool the S2 layer output over different size max pool filters.

The convolutional filters in the HMAX model use various kernel sizes (8, 10, 12, 14, 16, 18, 20, 22), as the network is designed to pool over different spatial scales in parallel.

#### Conv2d Model [27]

1. Conv2d(3, 64, (3, 3), (1, 1), 1)
2. ReLU()
3. Conv2d(64, 64, (3, 3), (1, 1), 1)
4. ReLU()
5. MaxPool2d(2, 2, 0)
6. Conv2d(64, 128, (3, 3), (1, 1), 1)
7. ReLU()
8. Conv2d(128, 128, (3, 3), (1, 1), 1)
9. ReLU()
10. MaxPool2d(2, 2, 0)
11. Linear(8192, 256)
12. ReLU()
13. Linear(256, 10)

### Experiment Details and Hyperparameters

#### VGG16 Networks

The Single Flexible Layer VGG16 networks use stochastic gradient descent with a learning rate = 0.001, momentum = 0.9 and weight decay = 0.006 for all parameters, except the Flexible Layer threshold which has weight decay = 0. A learning rate schedule is implemented with step size = 20 and gamma (the decay factor) = 0.7. The loss function is cross entropy loss. The networks are trained on the CIFAR-10 dataset for 150 epochs with batch size = 100. All images from the CIFAR-10 dataset are normalised and the images from the training set are also subject to random horizontal flip, with probability 0.5.

The All Flexible Layer networks are trained as above, but with learning rate = 0.01 and for 500 epochs on the CIFAR-10 dataset.

The Single Flexible Layer Networks trained on a subset of ImageNet use learning rate = 0.001, and batch size 50 due to memory constraints. The scheduler and other parameters are the same as above.

#### Small Network and HMAX Network

For the Small Network and the HMAX network, stochastic gradient descent with the same hyperparameters and schedule as used for the VGG16 networks is implemented. These networks are trained for 150 epochs on the FMNIST dataset. The batch size is always 100, and images from the FMNIST dataset are normalised.

### Datasets

#### Fashion-MNIST

Fashion-MNIST (FMNIST) [21] is a common benchmark dataset consisting of 60,000 images of clothing belonging to 10 classes. It was developed to provide more realism and difficulty to the benchmark MNIST dataset, performance on which has largely saturated in modern deep learning models. Each example is a 28x28 greyscale image. Validation is performed on an additional test set of 10,000 examples.

#### CIFAR-10

CIFAR-10 dataset [19] is an established computer-vision dataset used for object recognition. The training set consists of 50,000 32x32 colour images (3 colour channels) of 10 object classes. The validation set consists of an additional 10,000 examples.

#### Imagenette

Imagenette [20] is a subset of ImageNet [43], composed of the following 10 classes: tench, English springer, cassette player, chain saw, church, French horn, garbage truck, gas pump, golf ball, parachute. Images are cropped to 254x254 pixels for training and validation.

### Distinct activation patterns drive classification in flexible networks

In Fig. S1, a distinct, input-agnostic pattern can be observed in Flexible Layer’s feature maps in VGG16 network. Activations cluster in a “blocky” pattern which can appear as early as block 2.

**Fig S1.**
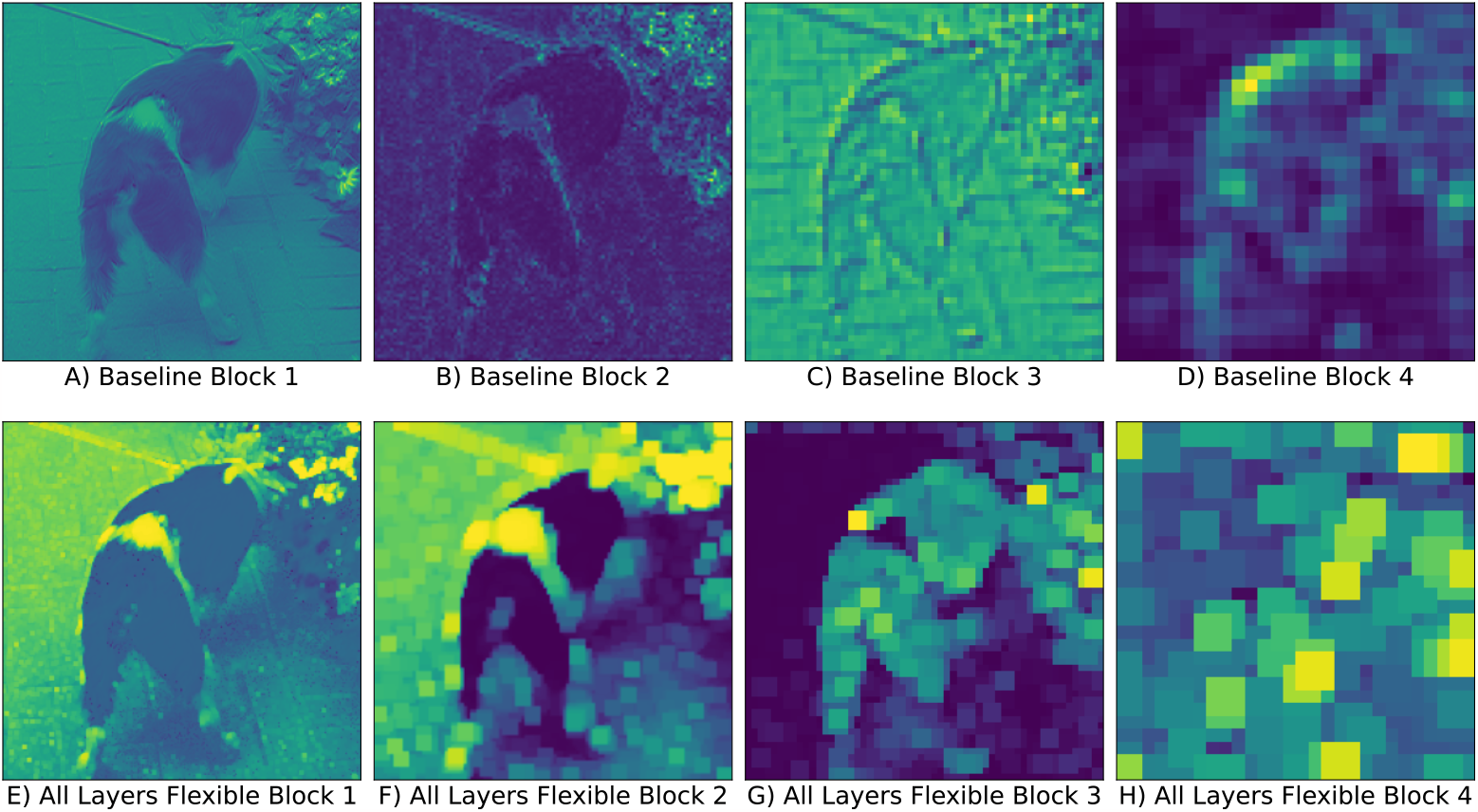
Example activation maps of consecutive layers in the network. (**A-D**) Activation maps of the convolution layer for an example image in block 1-4 respectively in the baseline VGG16 network. **(E-H)** Activation maps for an example image in a VGG16 network were all convolution layers have been replaced by flexible layers. Activations from block 1-4 respectively are shown.

## References

1. Hassabis D, Kumaran D, Summerfield C, Botvinick M. Neuroscience-Inspired Artificial Intelligence. Neuron. 2017;95(2):245–258. doi:10.1016/j.neuron.2017.06.011

2. Cox DD, Dean T. Neural networks and neuroscience-inspired computer vision; 2014.

3. Serre T, Oliva A, Poggio T. A feedforward architecture accounts for rapid categorization. Proceedings of the National Academy of Sciences. 2007;104(15):6424–6429. doi:10.1073/pnas.0700622104

4. Riesenhuber M, Poggio T. Hierarchical models of object recognition in cortex. Nature Neuroscience. 1999;2(11):1019–1025. doi:10.1038/14819

5. Hubel DH, Wiesel TN. Receptive fields, binocular interaction and functional architecture in the cat’s visual cortex. The Journal of physiology. 1962;160(1):106–154.

6. Vilankar KP, Field DJ. Selectivity, hyperselectivity, and the tuning of V1 neurons. Journal of Vision. 2017;17(9):9–9. doi:10.1167/17.9.9

7. Goris R, Op De Beeck H. Invariance in visual object recognition requires training: a computational argument. Frontiers in Neuroscience. 2010;3:12. doi:10.3389/neuro.01.012.2010

8. Poggio T, Mutch J, Anselmi F, Leibo JZ, Rosasco L, Tacchetti A. The computational magic of the ventral stream: sketch of a theory (and why some deep architectures work). Mit-Csail-Tr-2012-035. 2012;.

9. Kozlov AS, Gentner TQ. Central auditory neurons display flexible feature recombination functions. Journal of Neurophysiology. 2014;111(6):1183–1189. doi:10.1152/jn.00637.2013

10. Kouh M, Poggio T. A canonical neural circuit for cortical nonlinear operations. Neural Computation. 2008;20(6):1427–1451. doi:10.1162/neco.2008.02-07-466

11. Su J, Vargas DV, Sakurai K. One Pixel Attack for Fooling Deep Neural Networks. IEEE Transactions on Evolutionary Computation. 2019;23(5):828–841. doi:10.1109/TEVC.2019.2890858

12. Zhao P, Wang Y, Liu S, Lin X. An ADMM-based universal framework for adversarial attacks on deep neural networks. MM 2018 - Proceedings of the 2018 ACM Multimedia Conference. 2018; p. 1065–1073. doi:10.1145/3240508.3240639

13. Nguyen A, Yosinski J, Clune J. Deep neural networks are easily fooled: High confidence predictions for unrecognizable images. Proceedings of the IEEE Computer Society Conference on Computer Vision and Pattern Recognition. 2015;07-12-June-2015:427–436. doi:10.1109/CVPR.2015.7298640

14. Szegedy C, Zaremba W, Sutskever I, Bruna J, Erhan D, Goodfellow I, et al. Intriguing properties of neural networks. 2nd International Conference on Learning Representations, ICLR 2014 - Conference Track Proceedings. 2014; p. 1–10.

15. Elsayed GF, Papernot N, Shankar S, Kurakin A, Cheung B, Goodfellow I, et al. Adversarial examples that fool both computer vision and time-limited humans. Advances in Neural Information Processing Systems. 2018;2018-December:3910–3920.

16. Livnat A, Papadimitriou C, Pippenger N, Feldman MW. Sex, mixability, and modularity. Proceedings of the National Academy of Sciences of the United States of America. 2010;107(4):1452–1457. doi:10.1073/pnas.0910734106

17. Srivastava N, Hinton G, Krizhevsky A, Sutskever I, Salakhutdinov R. Dropout: A Simple Way to Prevent Neural Networks from Overfitting. Journal of Machine Learning Research. 2014;15(56):1929–1958.

18. Wang S, Wang X, Zhao P, Wen W, Kaeli D, Chin P, et al. Defensive dropout for hardening deep neural networks under adversarial attacks. IEEE/ACM International Conference on Computer-Aided Design, Digest of Technical Papers, ICCAD. 2018;doi:10.1145/3240765.3264699

19. Krizhevsky A. Learning Multiple Layers of Features from Tiny Images; 2009.

20. Imagenette; 2023. Available from: https://github.com/fastai/imagenette.

21. Xiao H, Rasul K, Vollgraf R. Fashion-MNIST: a Novel Image Dataset for Benchmarking Machine Learning Algorithms. arXiv. 2017;.

22. van Vliet M. PyTorch Implementation of HMAX; 2020. Available from: https://github.com/wmvanvliet/pytorch{_}hmax.

23. Serre T, Wolf L, Bileschi S, Riesenhuber M, Poggio T. Robust object recognition with cortex-like mechanisms. IEEE Transactions on Pattern Analysis and Machine Intelligence. 2007;29(3):411–426. doi:10.1109/TPAMI.2007.56

24. Ian J Goodfellow, Jonathon Shlens and Christian Szegedy. Explaining and Harnessing Adverserial ML. International Conference on Learning Representations (ICLR). 2015; p. 1–11.

25. Behnia F, Mirzaeian A, Sabokrou M, Manoj S, Mohsenin T, Khasawneh KN, et al. Code-bridged classifier (CBC): A low or negative overhead defense for making a CNN classifier robust against adversarial attacks. arXiv. 2020;(January).

26. Adversarial Example Generation — PyTorch Tutorials 2.0.1+cu117 documentation;. Available from: https://pytorch.org/tutorials/beginner/fgsm_tutorial.html.

27. Carlini N, Wagner D. Towards Evaluating the Robustness of Neural Networks. Proceedings - IEEE Symposium on Security and Privacy. 2017; p. 39–57. doi:10.1109/SP.2017.49

28. Zhang H, Yu Y, Jiao J, Xing EP, Ghaoui LE, Jordan MI. Theoretically Principled Trade-off between Robustness and Accuracy. International Conference on Machine Learning. 2019;.

29. Yu Y. TRADES (TRadeoff-inspired Adversarial DEfense via Surrogate-loss minimization). https://githubcom/yaodongyu/TRADES. 2023;.

30. Hendrycks D, Dietterich T. Benchmarking Neural Network Robustness to Common Corruptions and Perturbations. arXiv:190312261. 2019;.

31. Evans BD, Malhotra G, Bowers JS. Biological convolutions improve DNN robustness to noise and generalisation;148:96–110. doi:10.1016/j.neunet.2021.12.005

32. Mechler F, Ringach DL. On the classification of simple and complex cells. Vision Research. 2002;42(8):1017–1033. doi:10.1016/S0042-6989(02)00025-1

33. Jo J, Bengio Y. Measuring the tendency of CNNs to Learn Surface Statistical Regularities;. Available from: http://arxiv.org/abs/1711.11561.

34. Geirhos R, Rubisch P, Michaelis C, Bethge M, Wichmann FA, Brendel W. ImageNet-trained CNNs are biased towards texture; increasing shape bias improves accuracy and robustness;. Available from: http://arxiv.org/abs/1811.12231.

35. Malhotra G, Evans BD, Bowers JS. Hiding a plane with a pixel: examining shape-bias in CNNs and the benefit of building in biological constraints. Vision Research;174:57–68. doi:10.1016/j.visres.2020.04.013

36. Reddy MV, Banburski A, Pant N, Poggio T. Biologically Inspired Mechanisms for Adversarial Robustness; 2020.

37. Dapello J, Marques T, Schrimpf M, Geiger F, Cox DD, DiCarlo JJ. Simulating a Primary Visual Cortex at the Front of CNNs Improves Robustness to Image Perturbations. bioRxiv. 2020;doi:10.1101/2020.06.16.154542

38. Li Z, Ortega Caro J, Rusak E, Brendel W, Bethge M, Anselmi F, et al. Robust Deep Learning Object Recognition Models Rely on Low Frequency Information in Natural Images;19(3):e1010932. doi:10.1371/journal.pcbi.1010932

39. Jang H, Tong F. type [;]Available from: http://biorxiv.org/lookup/doi/10.1101/2023.07.29.551089.

40. Gaziv G, Lee MJ, DiCarlo JJ. Robustified ANNs Reveal Wormholes Between Human Category Percepts;. Available from: http://arxiv.org/abs/2308.06887.

41. wmvanvliet/pytorch hmax: Implementation of the HMAX model of vision in PyTorch;. Available from: https://github.com/wmvanvliet/pytorch{_{hmax.

42. Maxlab — HMAX Model;. Available from: https://maxlab.neuro.georgetown.edu/hmax.html.

43. Russakovsky O, Deng J, Su H, Krause J, Satheesh S, Ma S, et al. ImageNet Large Scale Visual Recognition Challenge. International Journal of Computer Vision (IJCV). 2015;115(3):211–252. doi:10.1007/s11263-015-0816-y

